# Retinotopic organization of head-orienting vectors in the pigeon ventral lateral geniculate nucleus

**DOI:** 10.64898/2026.10.03.756353

**Authors:** Daniel Severin, Bryan Reynaert, Tomas Vega-Zuniga, Juan Carlos Letelier, Gonzalo Marin, Harvey J. Karten, Jorge Mpodozis

**Affiliations:** Departamento de Biología, Facultad de Ciencias, Universidad de Chile, Santiago, 7800003, Chile; Mind/Brain Institute, Johns Hopkins University, Baltimore, MD, United States; Facultad de Medicina y Salud, Universidad Finis Terrae, Santiago, 7501015, Chile; Department of Neurosciences, University of California San Diego, San Diego, California, USA

## Abstract

The ventral lateral geniculate nucleus (GLv) is a conserved retinorecipient structure positioned between visual and premotor systems, yet its role in visuomotor transformations remains poorly understood. Here, we examined whether the pigeon GLv contains a spatially organized motor representation linking visual-field locations to orienting movements. Anatomical tracing identified descending projections from neurons in the GLv inner lamina (GLv-li) through pretectal and tegmental regions toward pontine targets. GLv-li projection neurons are GABAergic, yet electrical microstimulation of GLv-li in awake pigeons evoked rapid head movements followed by smaller eye movements. Each stimulation locus produced a reproducible characteristic motor vector whose direction and duration were largely insensitive to stimulation parameters, whereas movement amplitude and velocity approached site-specific plateaus. Movements initiated from different head positions were better explained by repetition of a fixed displacement than by convergence toward a common spatial goal. Critically, motor-vector direction was systematically related to the direction of the visual receptive field represented at the same GLv locus, while motor-vector amplitude covaried with receptive-field eccentricity. These defining features persisted after optic tectum ablation. Together, these findings reveal a retinotopically-organized map of fixed-vector head movements in the avian GLv and identify a tectum-independent route through which visual spatial representations can access descending orienting circuitry.

**SIGNIFICANCE:** Orienting toward an object requires the brain to translate visual locations into movements. This transformation is classically associated with the optic tectum, or superior colliculus in mammals, but how other visual structures contribute remains less clear. We show that the pigeon ventral lateral geniculate nucleus contains a map of head movements aligned with its established visual map. Head movements evoked by stimulating this nucleus persist after tectal removal, revealing an additional route through which visual spatial representations can access orienting circuits. These findings establish a spatially organized motor role for a conserved inhibitory visual thalamic structure and broaden our understanding of how the brain controls gaze.

## INTRODUCTION

The ventral lateral geniculate nucleus—GLv in birds and reptiles, vLGN in mammals—is a conserved component of the retinorecipient lateral geniculate complex in amniotes, yet it remains understudied relative to the canonical tectofugal and thalamofugal visual pathways (Niimi et al., 1963; Butler and Northcutt, 1971; Crossland and Uchwat, 1979). Prior studies have linked the GLv/vLGN to visual discrimination, circadian regulation, pupillary and optokinetic responses, and visuomotor control (Büttner and Fuchs, 1973; Pateromichelakis, 1979; Maturana and Varela, 1982; Hughes and Chi, 1983; Pickard and Turek, 1983; Jones, 1985; Gioanni et al., 1991; Guiloff, 1991; Harrington, 1997). Recent work in mice has shown that the vLGN regulates superior-colliculus-dependent visually evoked behavior (Fratzl et al., 2021), shapes visuospatial processing in the superior colliculus (Li et al., 2023), and coordinates visual signals with action-related motor signals during visuomotor behavior (Vega-Zuniga et al., 2025). Together, these findings place the vLGN at the interface between visual processing and action, but whether this visuomotor relationship has an internal spatial organization remains unknown.

Comparative anatomy situates the avian GLv within a broader ventral geniculate system whose organization differs from that of mammals. Whereas in mammals the ventral geniculate system comprises several subdivisions clustered within a relatively compact complex, in birds comparable components are more spatially segregated among distinct nuclei, including the GLv (Vega-Zuniga et al., 2014). This spatial segregation allows the GLv to be targeted as a discrete component of the broader system. The GLv is nested in the optic tract in the ventrolateral diencephalon and contains two main layers: the lamina interna (GLv-li), which contains projection neurons, and the neuropil (GLv-ne), a retinorecipient domain where retinal terminals contact GLv-li dendrites (Crossland and Uchwat, 1979; Guiloff et al., 1987; Guiloff, 1991; Tömböl et al., 2004). The retinal projection to GLv-ne is topographically organized and aligned with afferents from the optic tectum and visual Wulst (Karten et al., 1973; Crossland and Uchwat, 1979; Miceli et al., 1987; Shimizu et al., 1994; Vega-Zuniga et al., 2014). Both structures are linked to orienting behavior: stimulation of the optic tectum evokes head-based gaze shifts in barn owls (du Lac and Knudsen, 1990), whereas stimulation of the visual Wulst evokes topographically organized head movements in pigeons (Cohen and Pitts, 1967).

On the efferent side, GLv-li neurons are GABAergic (Güntürkün and Karten, 1991; Veenman and Reiner, 1994), and their axons follow a descending course to the nucleus pontis medialis (PM) in pigeons (Marín et al., 2001). In addition, detailed anatomical studies have shown that the GLv neurons send axon collaterals to ventrothalamic, pretectal, tectal, and tegmental targets (Vega-Zuniga et al., 2016; Vega-Zuniga et al., 2018). The PM-directed pathway places the GLv within a pontine route conveying visual information to folia VI–VIII of the avian oculomotor cerebellum (Wylie et al., 2018), and the tectal and tegmental descending pathways provide a plausible anatomical route for influencing gaze orientation (Masino and Knudsen, 1993; Scudder et al., 2002; Sparks, 2002; Yang et al., 2008; Wylie et al., 2018). We therefore tested whether electrical microstimulation of GLv-li sites evokes head and eye movements, and whether these movements are spatially related to the visual receptive fields at the stimulated sites.

Our results show that electrical microstimulation of GLv-li in pigeons evokes stereotyped, retinotopically coded head movements, with each stimulation site producing a characteristic motor vector systematically related to its visual receptive field. Crucially, these responses persisted after optic tectum ablation, indicating that GLv-li activation recruits a tegmental pathway capable of generating retinotopically organized head-movement vectors independently of the optic tectum.

## RESULTS

### Descending GLv efferents to PM arise from neurons in the inner lamina

We first characterized the descending GLv pathway to PM and identified its laminar origin. An anterograde PHA-L injection in the medial GLv labeled two predominantly ipsilateral efferent streams, coursing dorsomedially toward zona incerta (Zi) and dorsolaterally through nucleus pretectalis precommissuralis (PPC) around nucleus rotundus (Rt) (Fig. 1A–B). More caudally, labeled axons traversed the pretectum, turned medially and ventrally through the mesencephalic tegmentum, and continued to the ventral pons, where labeling was concentrated in PM (Fig. 1C–E). We then injected the retrograde tracer cholera toxin subunit β (CTB) into PM to determine the laminar origin of this pathway (Fig. 1F). Retrogradely labeled somata were concentrated in the ipsilateral GLv-li, with little somatic labeling in GLv-ne. These PM-projecting neurons extended prominent dendritic arborizations into the retinorecipient GLv-ne (Fig. 1G–H). Together, these tracing experiments identify GLv-li neurons as the source of a descending pathway that courses through pretectal and tegmental regions to PM.

**Fig. 1.**
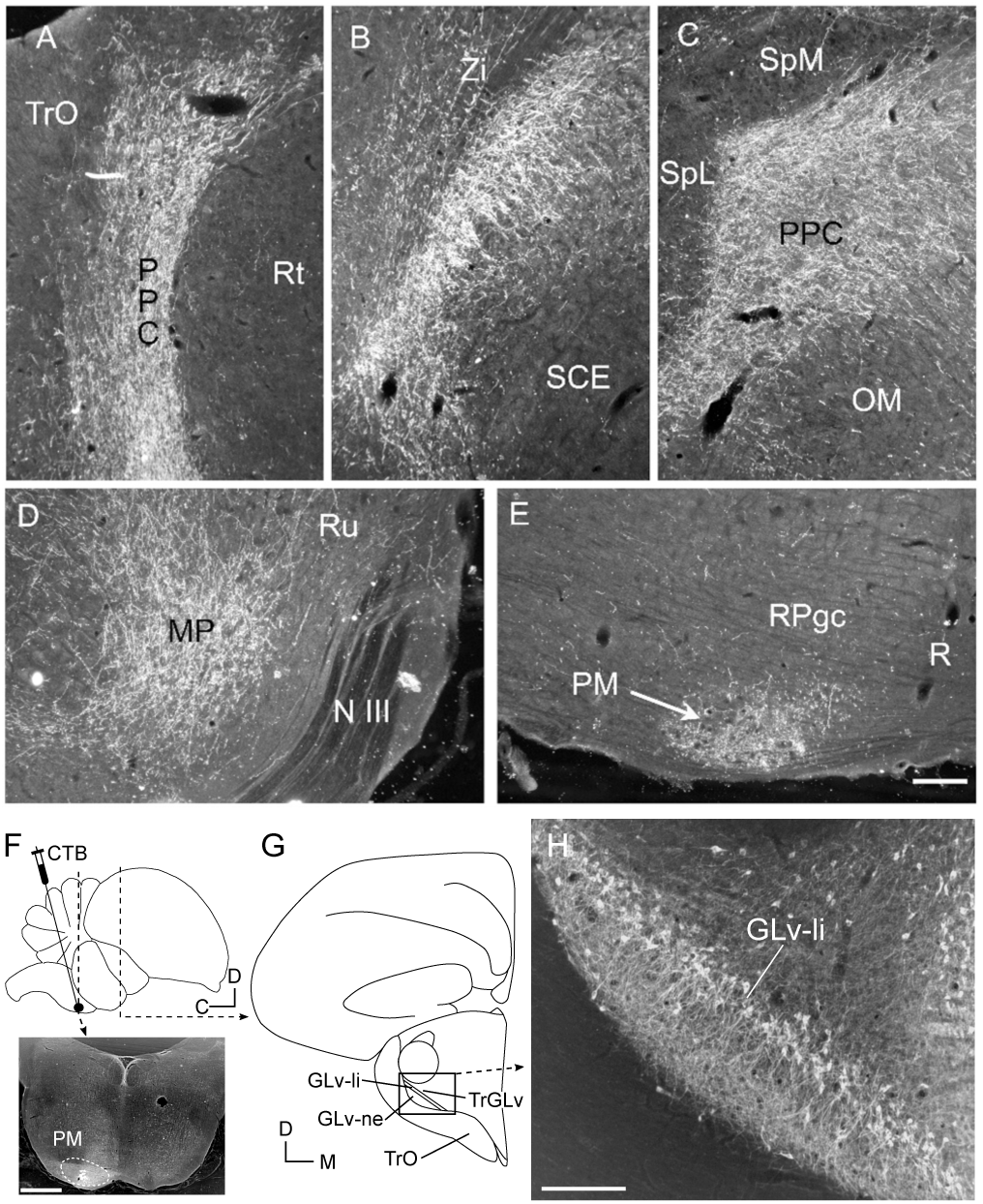
GLv descending efferents arise from neurons in the inner lamina. (A–E) Coronal sections showing the trajectory of anterogradely labeled GLv efferents after a PHA-L injection in medial GLv. Labeled fibers course dorsomedially toward Zi and dorsolaterally through PPC, then descend through the mesencephalic tegmentum toward PM. (F) Top, schematic of the CTB injection into PM; dashed lines indicate the section levels shown below and in G–H. Bottom, coronal section showing the CTB injection site in PM. (G) Schematic of the GLv at the level shown in H. (H) Coronal section showing PM-projecting GLv neurons retrogradely labeled from PM. Labeled somata are concentrated in GLv-li, with labeled processes extending into GLv-ne. Scale bars: 200 μm (A-E); 1 mm (F); 200 μm (H). Abbreviations: GLv-li, GLv lamina interna; GLv-ne, GLv neuropil; MP/PPN, nucleus pedunculopontine; N III, nervus oculomotorius; OM, nucleus nervi oculomotorii; PM, nucleus pontis medialis; PPC, nucleus pretectalis precommissuralis; R, nucleus raphes; RPgc, nucleus reticularis pontis caudalis pars gigantocellularis; Rt, nucleus rotundus; Ru, nucleus ruber; SCE, stratum cellulare externum; SpL, nucleus spiriformis lateralis; SpM, nucleus spiriformis medialis; TrGLv, GLv efferent tract; TrO, tractus opticus; Zi, zona incerta.

### GLv stimulation first evokes head movements, followed by eye movements

To determine whether activation of the GLv-li recruits head- and eye-related motor responses, we electrically stimulated visually responsive sites along vertical transects crossing the GLv, while monitoring eye movements and neck muscle twitches in anesthetized, head-fixed pigeons. Initial transects extending from the telencephalon to the optic tract revealed small contralateral eye movements, with the largest responses evoked from the GLv-ne and from the region where GLv-li efferent axons emerge (Fig. 2A–B; N = 3 transects in 3 animals). Small eye movements were also detected at two sites dorsal to the GLv, possibly reflecting current spread into the nucleus. At GLv sites, neck twitches were elicited at stimulation intensities lower than those required to evoke eye movements. To localize stimulation sites relative to GLv layers, we used visually evoked potentials and current source density (CSD) analysis as physiological landmarks. The GLv displayed a characteristic CSD profile, with a sink in the GLv-ne and a source in the GLv-li (Fig. 2C–F). After aligning stimulation sites to the center of the GLv-li, electrically evoked neck responses varied significantly with depth (linear mixed-effects model, depth as a fixed effect and transect as a random effect: χ²(16) = 93.45, P < 0.0001). The response profile showed a broad maximum centered approximately 200 μm dorsal to the GLv-li, overlapping the GLv efferent tract (Fig. 2G–H). Stimulation near the GLv-li also evoked small bilateral eye movements, with horizontal and vertical components ≤ 6° and variable directions across sites, including conjugate and vergence movements (Supplementary table 1). Thus, head-fixed experiments showed that GLv stimulation can recruit both ocular and cervical motor outputs, but could not determine whether the neck twitches corresponded to overt head movements and whether head movements preceded the evoked eye movements.

**Fig. 2.**
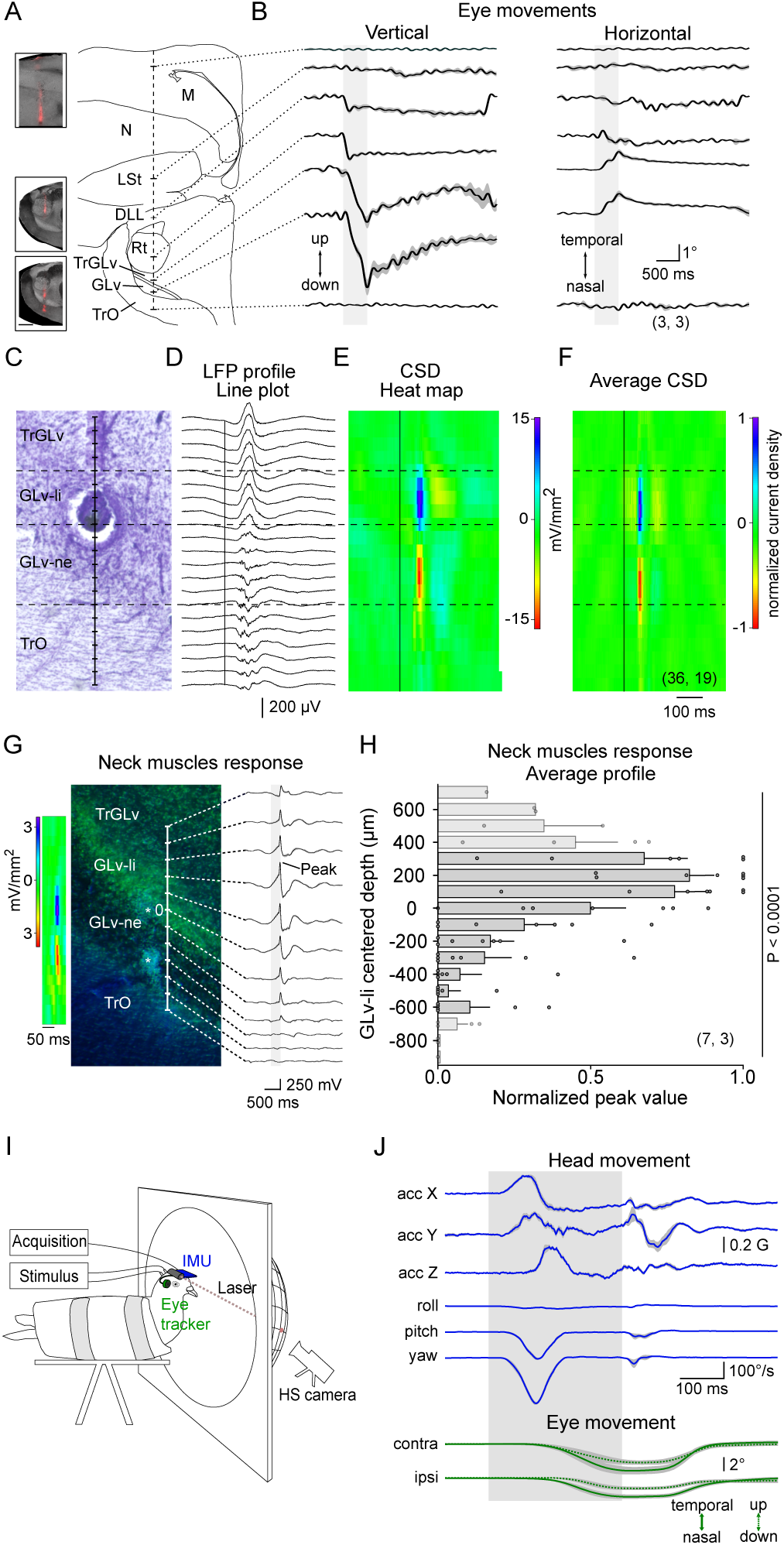
GLv stimulation recruits head-related motor output and eye movements. (A–B) Electrical stimulation was delivered along vertical transects from the visual Wulst to the optic tract. (A) Representative reconstructed transect with dye marks. (B) Average of vertical and horizontal displacements of the contralateral eye from corresponding sites across three transects in three animals; each site averaged from 10 trials. Corresponding sites were aligned using dye marks. Stimulation: cathodic monophasic, 300 μA, 300 Hz, 300 μs, 300 ms train. (C) Histological section from an example GLv transect with an electrolytic lesion in GLv-li. (D) Flash-evoked LFPs recorded at 21 depths spaced 50 μm apart. (E) CSD heat map derived from the LFP profile in D; red indicates sinks and blue indicates sources. (F) Average CSD profile showing a sink in GLv-ne and a source in GLv-li. Thin vertical lines indicate stimulus onset. Temporal scale for LFP and CSD traces is shown at bottom right. (G–H) Localization of neck responses relative to GLv layers. (G) Representative CSD profile, histological section, and neck-muscle responses evoked at 100 μm intervals. Asterisks indicate electrolytic lesions in GLv-li and GLv-ne. Stimulation: cathode-first, 150 μA, 300 Hz, 300 μs, 300 ms train. (H) Normalized neck responses varied significantly with depth (linear mixed-effects model, transect as random intercept: χ²(16) = 93.45, P < 0.0001). (I) Awake head-free preparation for simultaneous eye and head-motion tracking. (J) Example GLv-li–evoked head and bilateral eye movements shown as average traces ± SEM from 10 trials. Note that head movements precede eye movements. Stimulation: cathode-first, 300 μA, 300 Hz, 300 μs, 300 ms train. Gray bars indicate stimulus duration in B, G, and J. Sample sizes are shown as (transects, animals). Abbreviations: DLL, nucleus dorsolateralis anterior thalami pars lateralis; LSt, lateral striatum; M, mesopallium; N, nidopallium; Rt, nucleus rotundus; TrGLv, efferent tract of GLv; TrO, tractus opticus. Scale bar, 1 mm (A inset).

Because stimulation of gaze-control structures can recruit neck muscles before overt gaze shifts, neck activity alone cannot determine whether the response becomes a head movement or how it is timed relative to eye movements (Guitton et al., 1980; Roucoux et al., 1980; Corneil et al., 2002a,b; Elsley et al., 2007). We therefore tested GLv-li stimulation in awake, head-free animals. In these experiments, stimulating electrodes were implanted in the GLv-li, eye movements were measured with a head-mounted eye-tracking system, and head motion was recorded with an inertial measurement unit (IMU) calibrated by laser projection onto a campimeter (Fig. 2I and Supplementary fig. 1). Electrical stimulation of the GLv-li elicited rapid, saccadic head movements involving translations and rotations along yaw, pitch, and roll axes. These head movements were consistent across trials and were followed by smaller eye movements ipsilateral and contralateral to the stimulation site. Across implanted sites, the threshold for evoking head movements was lower than that for evoking eye movements, and head movement latencies were consistently shorter (Fig. 2J and Supplementary table 1). Together, these results indicate that GLv-li stimulation engages a gaze-related motor pathway in which head movements are recruited first, followed by smaller eye movements. We next asked whether the direction and amplitude of these head responses were topographically organized.

### Stereotyped head movements define a characteristic motor vector for each GLv locus

To address this, analyzed the initial head response evoked by GLv-li stimulation. For these experiments, the head was returned to the same standard initial position before each trial, allowing the calibrated IMU to reconstruct the full movement trajectory. Across six stimulation loci in five animals, projected beak trajectories were highly consistent across trials but differed across stimulation sites (Fig. 3A). The corresponding angular position and velocity profiles showed similarly stereotyped temporal dynamics (Fig. 3B), and a multidimensional summary of movement parameters suggested that each site was associated with a distinct kinematic profile (Fig. 3C). These observations prompted us to quantify which features of the evoked head movements were determined by stimulation parameters and which were determined by the stimulated GLv locus.

**Fig. 3.**
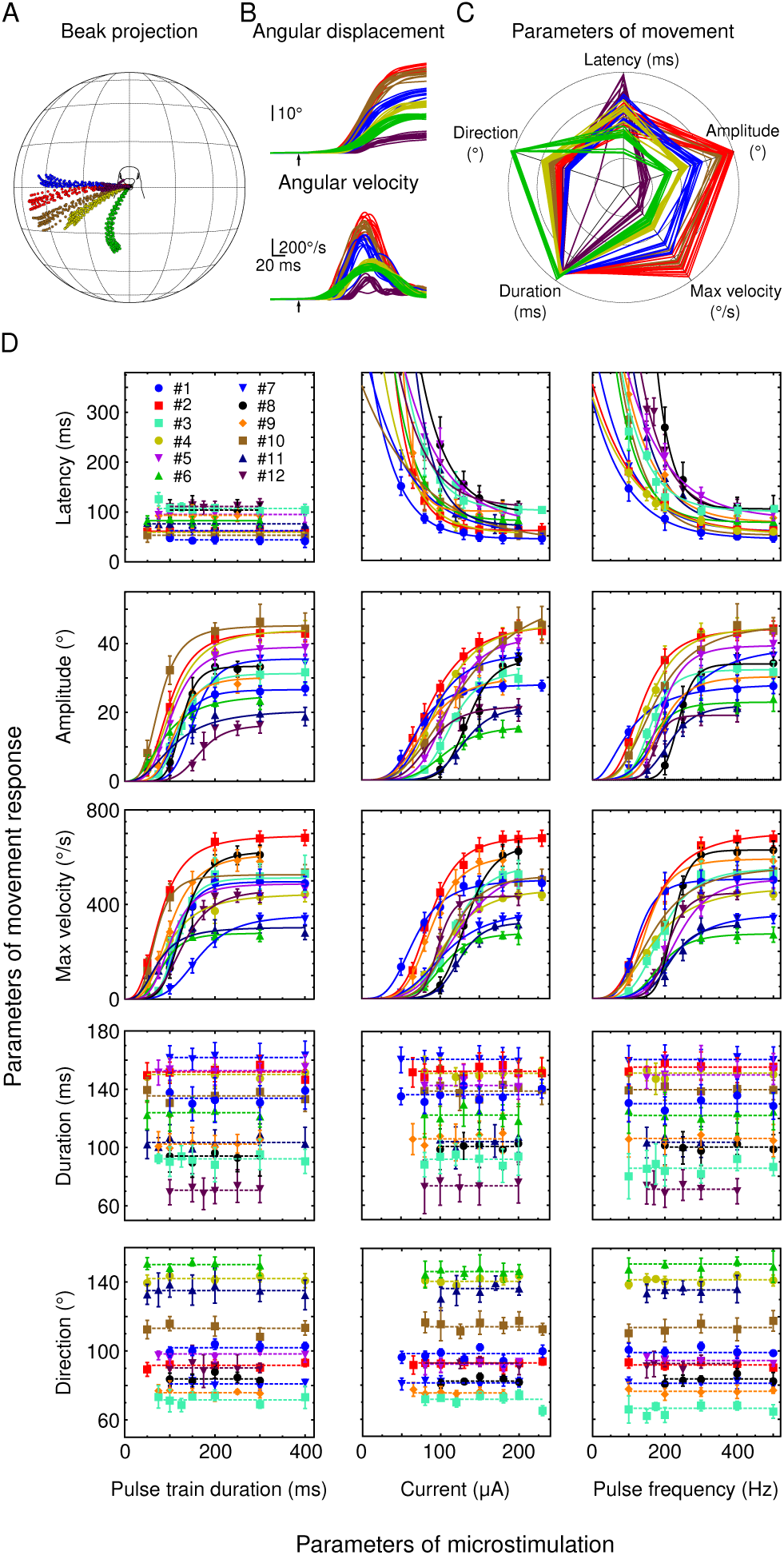
GLv loci generate reproducible, site-specific head-movement vectors. (A) Beak projections showing trial-by-trial reproducibility of electrically evoked head movements from six GLv loci; colors indicate stimulation loci, 10 trials per locus. (B) Angular displacement and velocity traces for the movements shown in A. The six example loci shown in A–C correspond to sites #2, #4, #6, #7, #10, and #12 in D; for A–C, stimulation parameters were fixed within each locus but could differ across loci. (C) Radar plot of individual-trial kinematic parameters for the six loci shown in A–B. Axes are scaled independently with zero at the center, illustrating clustering of trials by stimulation locus. (D) Effects of pulse-train duration, current, and pulse frequency on head-movement kinematics across 12 GLv sites in 8 pigeons. Symbols show mean ± SD from 10 trials. Lines show linear, exponential, or sigmoidal fits; dashed lines indicate nonsignificant fits (extra sum-of-squares F test, P > 0.05).

We systematically varied current amplitude, train duration, and stimulation frequency across 12 sites in 8 pigeons to distinguish stimulation-dependent from site-dependent features (Fig. 3D). Within the tested ranges, movement direction and duration were not significantly affected by any stimulation parameter (Extra sum-of-squares F test, hereafter F test, P > 0.05). Latency was unaffected by train duration but decreased exponentially with current amplitude or stimulation frequency in all but two site–parameter fits (F tests, P < 0.05; exponential fit R^2^ > 0.9386 for significant fits). Movement amplitude and maximum velocity increased sigmoidally with stimulation strength in all but three site–parameter fits (F tests, P < 0.05; sigmoidal fit R^2^ > 0.9218 for significant fits). However, the plateau values for movement amplitude and maximum velocity differed significantly across stimulation sites, even under identical stimulation conditions (F test, P < 0.0001). Thus, under saturating stimulation, each GLv locus was associated with a distinct combination of movement direction, duration, amplitude, and maximum velocity. We define this site-specific movement output as the characteristic motor vector (CMV), indicating that GLv-evoked head movements are topographically coded.

### GLv stimulation evokes fixed-vector head movements consistent with retinally organized motor outputs

Because the dependence of an evoked orienting movement on initial head position can distinguish different spatial codes, we compared two models: a fixed-vector model, in which stimulation evokes the same displacement from different starting positions, and a goal-directed model, in which movements from different starting positions converge toward a common endpoint. In a separate set of six pigeons, we tested six GLv sites while varying the initial head position, unlike the standardized starting position used for Fig. 3. Because the IMU calibration applied only to the standard position, these movements were measured from the laser projection with a high-speed camera, extracting initial and final head positions (Fig. 4A–F). Across sites, movements initiated from different head positions showed similar directions and displacements within each site, qualitatively consistent with a fixed-vector organization. Changing the initial head position produced relatively modest changes in movement direction and amplitude, although these effects varied across sites (Fig. 4G–J). We then compared the predictive accuracy of fixed-vector and goal-directed models using leave-one-trial-out cross-validation. The fixed-vector model yielded lower RMS endpoint prediction errors at all six sites (paired Wilcoxon signed-rank test, P = 0.0312; Fig. 4K), indicating that GLv-evoked head movements are better described by site-specific motor vectors than by movements toward spatial goals.

**Fig. 4.**
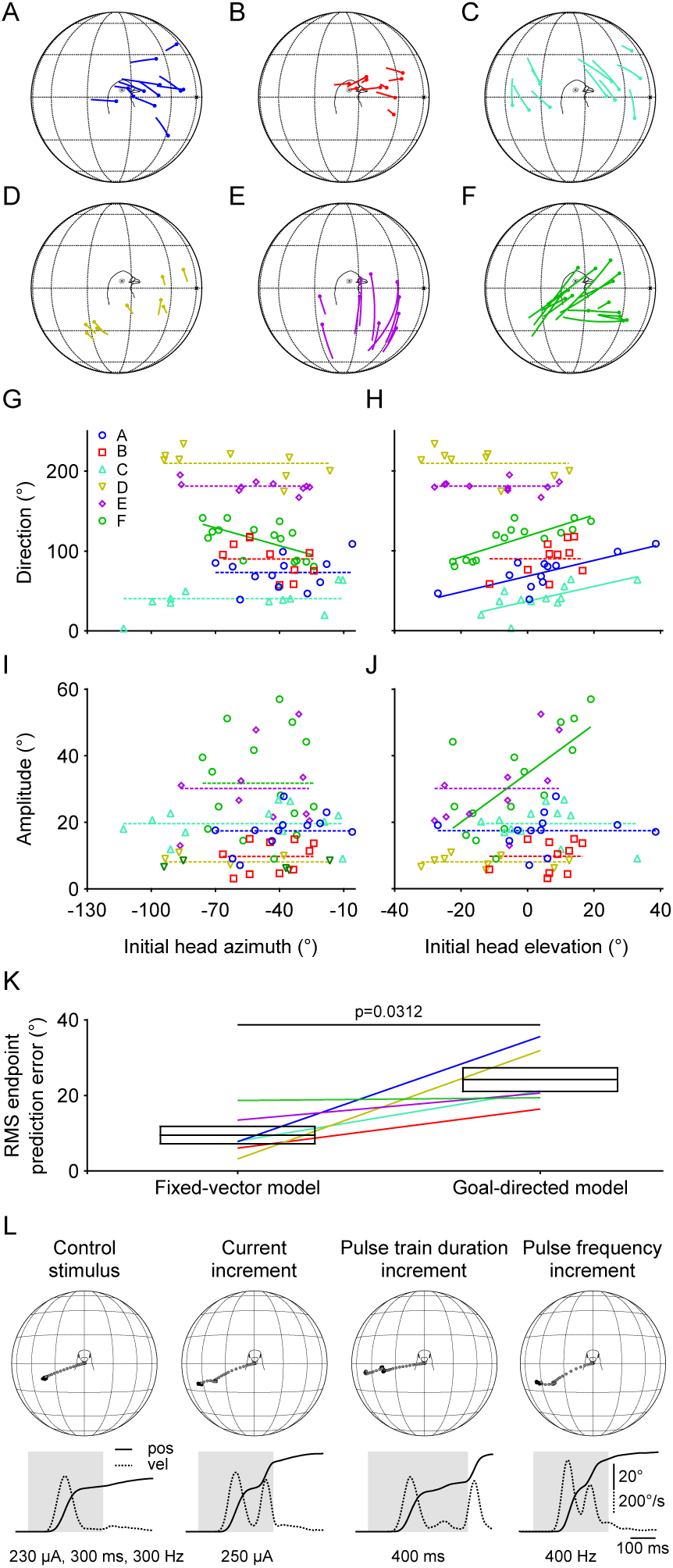
GLv stimulation evokes fixed-vector head movements across initial head positions. (A–F) Head movement vectors evoked from six GLv sites using stimulation parameters that elicited each site’s characteristic motor vector. Vectors show initial and final head positions from 9–15 movements per site. (G–J) Relationships between initial head azimuth or elevation and movement direction or amplitude for the six sites shown in A–F. Lines show the corresponding fits; direction and amplitude are defined as in Fig. 3. (K) RMS endpoint prediction errors for fixed-vector and goal-directed models for the same six sites shown in A–J, using leave-one-trial-out cross-validation; lines connect the same stimulation site. Errors were lower for the fixed-vector model (paired Wilcoxon signed-rank test, P = 0.0312). Boxes show mean ± SEM. (L) Beak trajectories and corresponding angular position (pos) and velocity (vel) traces during increasing current, pulse-train duration, or pulse frequency. Stronger stimulation produced stepwise repetitions of the site-specific vector. Gray bars indicate stimulus duration.

We also tested whether this vector organization persisted when stronger or longer stimulation recruited additional saccades. Once stimulation exceeded the threshold for the initial CMV, increasing current, pulse-train duration, or pulse frequency often evoked a second or additional saccades after a brief fixation (Fig. 4L). These subsequent saccades began from the new head position but usually preserved the same direction, producing a staircase-like sequence in which the site-specific vector was repeated as the head was displaced. Together, the lower cross-validated prediction error of the fixed-vector model and the preservation of direction across successive saccades support a fixed-vector rather than goal-directed organization. Because this organization is consistent with a retinotopically-organized motor output, we next asked whether the characteristic motor vector evoked at each GLv locus corresponded spatially to the receptive field represented at that site.

### Characteristic motor vectors are systematically related to receptive fields in the GLv

To test for spatial correspondence between the sensory and motor maps in the GLv, we mapped the visual receptive field (RF) location at different recording sites in the GLv and compared them with the CMV evoked by electrical stimulation of those sites. RFs were mapped by measuring visually evoked potential (VEP) amplitudes elicited by a 4° circular spot of collimated light flashed at different visual-field locations (Fig. 5A). Because frontal and eccentric visual fields differ in their representation within the pigeon GLv, we compared frontal RFs, defined as those within 60° of the beak projection, with eccentric RFs. Frontal RFs were smaller than eccentric RFs (Fig. 5B–C; frontal, 3.57 ± 0.34°, N = 9 loci, 7 animals; eccentric, 5.47 ± 0.81°, N = 8 loci, 7 animals; Lognormal t test: t = 2.42, P = 0.0288). We then compared the direction and amplitude of each CMV with the direction of the corresponding RF center relative to the beak projection and its eccentricity across 25 sites in 15 animals (Fig. 5D–F). CMV direction was positively correlated with RF-center direction in both frontal and eccentric regions (Fig. 5E; frontal, r = 0.8882, P < 0.0001; eccentric, r = 0.6604, P = 0.0194). The regression slopes differed between groups (F[1,21] = 4.475, P = 0.0465), with the slope for the frontal RFs not different from unity (F[1,11] = 0.4895, P = 0.4987), whereas the eccentric RFs’ slope was (F[1,10] = 11.78, P = 0.0064). CMV amplitude was also correlated with RF-center eccentricity in both groups (Fig. 5F; frontal, r = 0.7999, P = 0.0010; eccentric, r = 0.7227, P = 0.0079). The regression intercepts differed between groups (F[1,22] = 10.21, P = 0.0042), whereas the slopes did not (F[1,21] = 1.216, P = 0.2826). Thus, CMV direction followed the corresponding RF-center direction, most closely in the frontal visual field, while CMV amplitude covaried with RF-center eccentricity rather than matching it directly.

**Fig. 5.**
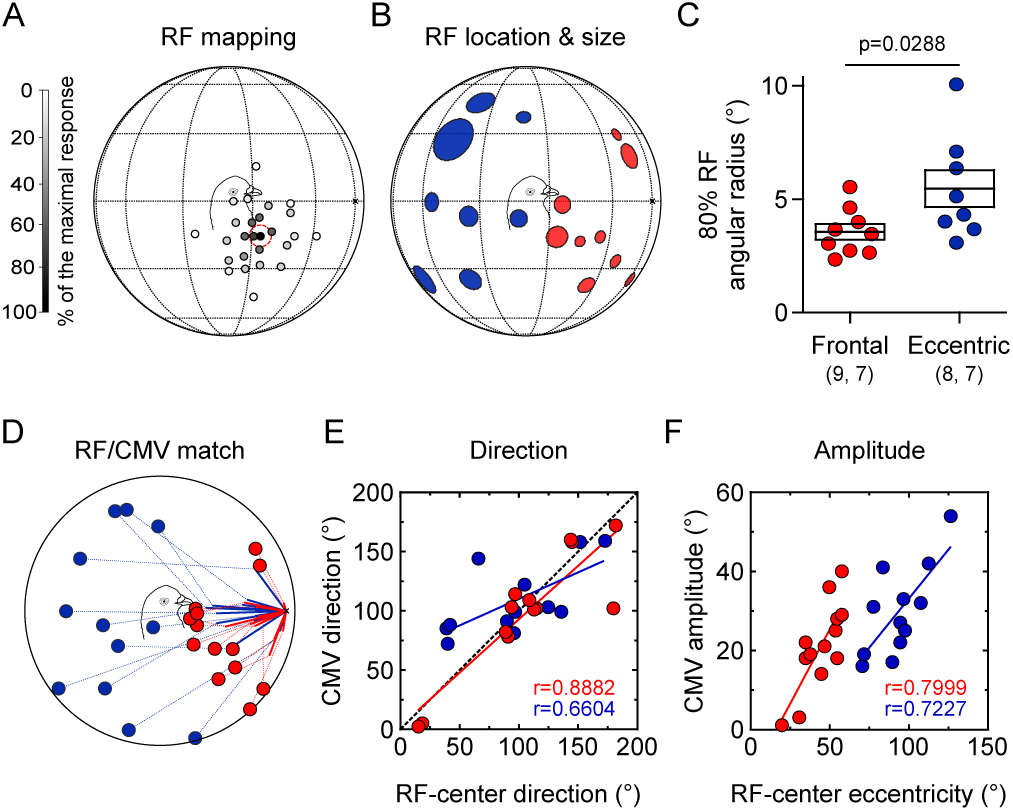
Correlation between GLv motor vectors and visual receptive fields. (A) Example receptive-field map from one GLv locus. Grayscale indicates VEP amplitude as percentage of the maximal response; filled circle marks the RF center. (B) RF locations and sizes across GLv sites. Red, frontal RFs; blue, eccentric RFs; x, eye–bill axis. Circle radius represents the 80% RF angular radius. (C) The 80% RF angular radius was smaller for frontal than eccentric RFs (lognormal t test, P = 0.0288). Sample sizes are shown as (sites, animals). Boxes show mean ± SEM. (D) RF centers and corresponding characteristic motor vectors (CMVs) for 25 GLv sites. Circles indicate RF centers; lines indicate CMVs; dotted lines connect each CMV endpoint to its corresponding RF. (E) Correlation between RF-center direction, measured from the beak projection, and CMV direction (Pearson correlation: frontal, r = 0.8882, P < 0.0001; eccentric, r = 0.6604, P = 0.0194); regression slopes differed between groups (P = 0.0465). (F) Correlation between RF-center eccentricity, measured from the beak projection, and CMV amplitude (frontal, r = 0.7999, P = 0.0010; eccentric, r = 0.7227, P = 0.0079); regression intercepts differed between groups (P = 0.0042), whereas slopes did not (P = 0.2826). (E–F) Red and blue lines indicate linear regressions for frontal and eccentric RFs.

### GLv-evoked characteristic motor vectors persist after optic tectum ablation

Because the GLv receives input from the tectum, a structure closely involved in head and eye movements, we next asked whether GLv-evoked CMVs required an intact optic tectum (OTe) or could result from retrograde or polysynaptic activation of tectal circuitry. To test this possibility, we examined GLv-evoked motor responses after OTe ablation.

Across post-ablation preparations, GLv stimulation continued to evoke eye and head responses. Eye movements persisted under both head-restrained conditions (n = 2 sites, 1 animal) and head-free conditions (n = 4 sites, 3 animals). Head movements also retained qualitatively similar stimulus–response relationships after ipsilateral OTe ablation (n = 4 sites, 2 animals), and the same qualitative result was observed at one site after bilateral OTe ablation. Complete paired pre- and post-ablation parametrization was obtained from two GLv sites in one animal and was used for quantitative comparison of CMV-defining parameters.

After ipsilateral OTe ablation, movement direction and duration remained independent of stimulation parameters and did not differ from pre-ablation values (two-way ANOVA, P > 0.05; Fig. 6; Supplementary fig. 2). Latency as a function of pulse-train duration was also unchanged (two-way ANOVA, P > 0.05). For current amplitude and pulse frequency, OTe ablation altered the input–output curves for latency, amplitude, and maximum velocity (two-way ANOVA, P < 0.0001), indicating a change in response sensitivity. However, the plateau values for movement amplitude and maximum velocity were preserved (F test, P > 0.05). Thus, although OTe ablation modified how some response parameters depended on stimulation strength, it did not alter the defining features of the CMV under saturating stimulation. These results support the idea that GLv-li activation recruits a tegmental pathway to generate retinotopically organized head-movement vectors independently of the optic tectum.

**Fig. 6.**
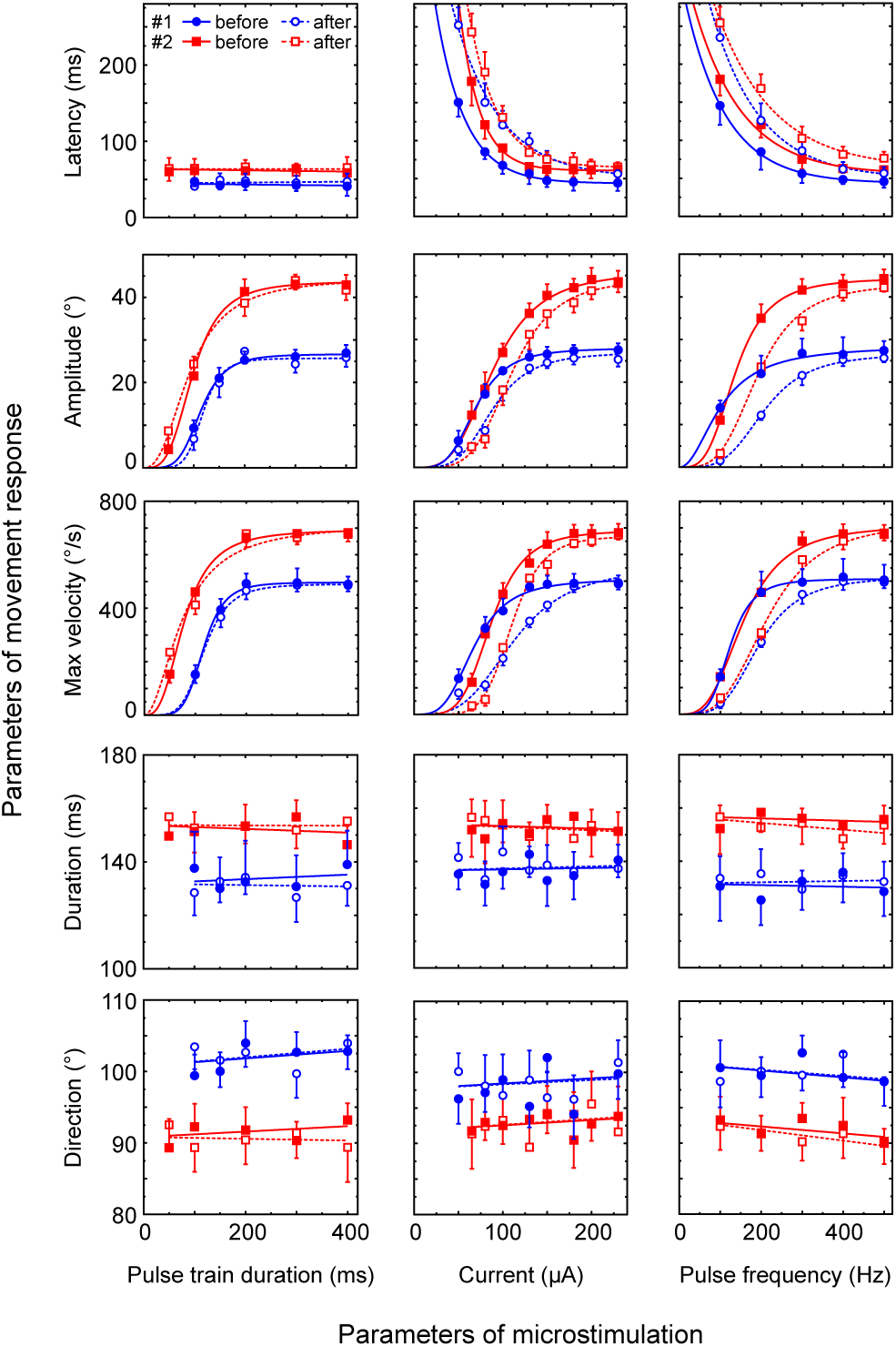
Optic tectum ablation preserves GLv-evoked motor-vector kinematics. Kinematic parameters of head movements evoked from two GLv sites in one pigeon before and after ablation of the OTe ipsilateral to the stimulated GLv. Closed symbols, pre-ablation; open symbols, post-ablation. Sites #1 and #2 are replotted from Fig. 3D, with the same plotting and fit conventions. Direction and duration were unchanged after ablation (two-way ANOVA, P > 0.05). Ablation altered current- and frequency-dependent responses for latency, amplitude, and maximum velocity (P < 0.0001), while amplitude and maximum-velocity plateaus were preserved (extra sum-of-squares F test, P > 0.05).

## DISCUSSION

The GLv has long been recognized as a conserved retinal target positioned between visual and premotor systems, but its contribution to visuomotor transformations has remained uncertain. Recent work in mice has shown that the vLGN regulates visual processing in the superior colliculus, visually guided behavioral selection, and action-related sensory signals (Fratzl et al., 2021; Li et al., 2023; Vega-Zuniga et al., 2025). Here, we showed that in the avian GLv the visuomotor transformation is conspicuously straightforward. GLv-li stimulation evoked rapid head movements that preceded smaller eye movements, and each stimulated locus produced a reproducible characteristic motor vector (CMV). Movement direction and duration were largely insensitive to stimulation parameters, whereas movement amplitude and velocity approached site-specific plateaus. Moreover, the evoked movements were better explained by a fixed-vector than a goal-directed model, and CMV direction and amplitude were systematically related to the receptive field represented at the same locus. Finally, the defining features of these vectors persisted after optic tectum ablation. Together, these findings identify the avian GLv as a spatially organized visuomotor structure that links local visual representations to site-specific vectors of head movement through a descending pathway that does not require an intact optic tectum.

This motor effect is notable because the projection neurons of GLv-li are GABAergic (Güntürkün and Karten, 1991; Veenman and Reiner, 1994). Long-range inhibition can initiate movement through disinhibitory circuit motifs. In the classic basal ganglia model, tonically active GABAergic neurons in substantia nigra pars reticulata (SNr) suppress superior-colliculus and other premotor targets, and inhibition of SNr releases these downstream circuits to permit orienting movements; this disinhibitory organization has support in monkeys, cats, rats, and lampreys (Hikosaka and Wurtz, 1983; Chevalier et al., 1985; Gelissen and Cools, 1986; Stephenson-Jones et al., 2012; Frost-Nylén et al., 2024). GLv output could operate through an analogous motif by inhibiting another inhibitory node and thereby releasing a descending premotor network. However, inhibition need not be merely permissive. Long-range GABAergic neurons can exert temporally precise control over distributed circuits (Roux and Buzsáki, 2015; Tremblay et al., 2016), and optogenetic activation of the inhibitory nigrocollicular pathway in mice can itself evoke orienting movements through post-inhibitory rebound in collicular output neurons (Villalobos and Basso, 2022). Thus, GLv inhibition could contribute actively to the timing and recruitment of head-orienting movements, rather than simply removing a brake. Identifying the postsynaptic targets of GLv-li axon terminals in the tegmentum and pons will be necessary to determine how these signals engage premotor networks coordinating head and eye movements.

The fixed-vector organization provides an important clue to the computation performed by this pathway. If a GLv locus specified a fixed goal in external space, stimulation from different starting head positions should drive movements toward a common endpoint. Instead, the same site generated approximately the same displacement from different initial positions, and stronger or longer stimulation could repeat that displacement in a staircase-like sequence. This organization resembles the vector maps described in the optic tectum and superior colliculus, where local activity specifies orienting movements of characteristic direction and amplitude (du Lac and Knudsen, 1990; Sparks, 2002; Gandhi and Katnani, 2011). In the GLv, these motor vectors were systematically linked to the underlying retinotopic representation: CMV direction pointed towards the RF location, while CMV amplitude covaried with RF-center eccentricity. Because the GLv receives a topographic tectogeniculate projection, electrical stimulation could in principle have recruited tectal neurons antidromically or through local polysynaptic loops. The persistence of CMVs after ipsilateral tectal ablation, together with qualitatively preserved responses after bilateral ablation in one case, argues that the GLv motor output is not simply inherited from acute recruitment of the tectal motor map. Tectal removal did alter some stimulus-response relationships, which could reflect loss of tectal drive to GLv or nonspecific changes associated with the chronic preparation, but the site-defining direction, duration, and saturated movement metrics were preserved. Combined with the descending GLv projection through the tegmentum to PM and other premotor regions, these results support a tectum-independent route by which retinotopically organized GLv activity can access head-orienting circuitry.

The present findings also fit a broader comparative view of the ventral geniculate system. Across amniotes, ventral geniculate structures retain strong retinal input and descending access to premotor and precerebellar systems, although their organization differs. In lizards, for example, the GLv receives retinotopically organized retinal and tectal inputs and projects to the red nucleus, providing an anatomical route for visuomotor control (Martínez-Marcos et al., 1999). In birds, the GLv is a discrete laminar nucleus, whereas comparable components of the mammalian ventral geniculate system are clustered within a more compact complex of subdivisions (Vega-Zuniga et al., 2014). A vLGN projection to medial pontine regions was described in mammals decades ago (Graybiel, 1974), and recent mouse studies show that GABAergic vLGN circuits regulate superior-colliculus-dependent behavioral output and visuospatial processing (Fratzl et al., 2021; Li et al., 2023). Vega-Zuniga et al. (2025) further identified the vLGN as a hub that combines visual and action-related signals and can drive corrective motor responses. The pigeon results extend this emerging visuomotor role by revealing a tighter relationship between visuospatial location and motor execution. At the same time, the differences in nuclear organization across amniotes argue against a strict one-to-one correspondence between the avian GLv and the entire mammalian vLGN complex.

A possible neuroethological correlate for the topography of the evoked head movements is suggested by the visual strategy pigeons use during pecking. Goodale described two head fixations for targets already located in the pigeon’s binocular field before the final peck: an earlier fixation (F1) associated with the decision to peck and a later, closer fixation (F2) used to specify target size, depth, and location (Goodale, 1983). Peripheral targets can also recruit an additional orienting head movement before pecking (Cesaretti et al., 1997). This sequence is compatible with the organization observed here. The larger, more variable vectors associated with eccentric GLv loci resemble the head movements leading to the earlier F1 fixation, which could bring a peripheral target toward the frontal field. Because CMV amplitude covaried with RF-center eccentricity without matching it, this transformation may support iterative rather than single-step target alignment, with successive movements progressively shifting a peripheral target into the frontal field. By contrast, the smaller receptive fields and closer RF–CMV directional correspondence of frontal GLv loci resemble the head movement leading to the later F2 fixation, when the target is already represented in the frontal/red-area field and finer spatial control is required. On occasion, the stimulation-evoked movement sequence culminated in a peck-like motion. The strong tectal and Wulst inputs to GLv could allow this retinal map to be modulated by ongoing visuomotor context during approach.

The tectal and Wulst inputs to GLv may shape when and where its retinotopic motor output is recruited. In pigeons, the Wulst is preferentially engaged by distant lateral vision and contributes to large-scale spatial representation (Clark and Colombo, 2022). Wulst input could also provide top-down, experience-dependent context for whether a peripheral object is worth orienting toward and bringing into the frontal field. The optic tectum, in turn, is embedded in an isthmic circuit in which focal re-entrant signals from the parvocellular isthmic nucleus enhance selected tectal locations (Marín et al., 2005), while wide-field inhibitory interactions generate competition among spatial representations (Marín et al., 2007; Mysore et al., 2010; Mysore and Knudsen, 2011; Marín et al., 2012). This circuit could bias which of several simultaneously active GLv loci drives the next orienting movement through the tecto-GLv projection, which is topographic and cholinergic (Vega-Zuniga et al., 2014), consistent with the ability of acetylcholine to bidirectionally modulate retinal transmission in chick GLv (Guo et al., 2010).

In summary, stimulation of the pigeon GLv-li reveals a retinotopically organized map of fixed-vector head movements. Individual GLv loci specify characteristic head displacements, and this organization persists after optic tectum removal, consistent with access to premotor tegmental and pontine structures through descending GLv-li projections. These findings identify the GLv as a spatially organized node capable of linking visual locations to head-orienting movements. Within the broader visuomotor network formed with tectal and telencephalic inputs, the GLv may exert inhibition-mediated control over how selected retinal locations gain access to descending orienting circuitry.

---

**Supplementary fig. 1. Calibration of the IMU for head-free recordings.** (A) Translational and rotational axes used to describe head motion. (B) Calibration workflow aligning the IMU-derived projection with the laser reference. IMU calibration was performed before recording; calibration parameters were accepted when the difference between IMU- and laser-derived projections was <2°.

**Supplementary fig. 2. Histological reconstruction of optic tectum ablation.** (A–B) Posterior and lateral views of the brain after ablation of the OTe ipsilateral to the stimulated GLv. (C) Coronal section series showing the reconstructed ablation; sections are spaced 180 μm apart. The anterior OTe remained intact, whereas the implanted GLv sites corresponded to intermediate and posterior OTe representations. Scale bar, 300 μm.

**Supplementary table 1. Eye and head movements evoked by GLv-li stimulation.** Threshold is the minimum current required to elicit a detectable movement. Latency is the shortest response onset observed across tested current levels. Amplitude is the maximum displacement observed across the tested stimulus range. Stimulus range: 50–300 μA, 30–400 ms pulse train, 300 Hz.

## METHODS

Detailed experimental procedures and analysis methods are provided in Supplementary Methods.

### Animal preparation and surgery

Experiments were conducted on 44 adult feral pigeons (300–420 g) of either sex. Animals were anesthetized with ketamine (75 mg/kg, i.m.) and xylazine (5 mg/kg, i.m.) and positioned stereotaxically according to Karten and Hodos (1967). Electrodes were implanted through a craniotomy overlying the GLv and secured with dental acrylic. In five animals, the optic tectum (OTe) was surgically ablated ipsilateral or bilateral to the stimulated GLv. Chronic recording sessions began 2 d after surgery. All procedures were approved by the Ethics Committee of the Science Faculty of the University of Chile and conformed to NIH guidelines for animal research.

### Electrical stimulation and physiological localization

GLv-li was stimulated with monopolar tungsten microelectrodes using constant-current pulse trains. Unless otherwise stated, pulses were biphasic and charge-balanced (300 μs cathodal followed by 300 μs anodal; 50–300 μA, 100–500 Hz, 30–400 ms). Stimulation parameters were varied to determine movement thresholds and stimulus–response functions. Visually evoked potentials (VEPs) and current source density (CSD) profiles were used to identify GLv-ne and GLv-li, and stimulation sites were subsequently confirmed histologically.

### Receptive-field mapping

Visual receptive fields (RFs) were mapped under anesthesia using a hemispherical screen centered on the right eye and inverse retinoscopy to establish retinal landmarks. VEPs were evoked by a 4° spot of collimated light presented at different visual-field locations. The RF center was defined as the location producing the maximal VEP, and RF size was quantified from the 80% isoline of a two-dimensional Gaussian fit.

### Movement recording and analysis

Eye movements were recorded at 60 Hz with a ViewPoint EyeTracker infrared video system (Arrington Research). Neck responses were measured with a Hall-effect sensor, and head movements in awake, head-free pigeons were recorded with a high-speed camera and an inertial measurement unit (IMU). IMU-derived trajectories were calibrated against a head-mounted laser projection onto a campimeter (Supplementary Fig. 1). For stimulus–response parametrization, the head was returned to the same standard initial position before each trial. Movement direction, amplitude, latency, duration, and maximum velocity were quantified from reconstructed trajectories. For the fixed-vector versus goal-directed analysis in Fig. 4A–K, movements were instead initiated from different head positions and quantified from the laser projection by extracting initial and final positions; we then tested whether those movements were better explained by repetition of the same site-specific vector or by convergence toward a common endpoint.

### Neural tracing and histology

Anterograde GLv projections were labeled by iontophoretic injection of Phaseolus vulgaris leucoagglutinin (PHA-L) into medial GLv, and PM-projecting GLv neurons were labeled by pressure injection of cholera toxin subunit B (CTB) into PM. Brains were sectioned coronally and tracer labeling was visualized immunohistochemically. Stimulation sites were reconstructed from electrolytic lesions and, in some cases, fluorescent electrode-track labeling.

### Statistical analysis

Depth-dependent neck responses were analyzed with a linear mixed-effects model. Stimulus–response relationships were analyzed using exponential or sigmoidal fits and extra sum-of-squares F tests. Pre- and post-ablation responses were compared by two-way ANOVA. RF size was compared with a lognormal t test, RF–CMV relationships with Pearson correlations and linear regression, and fixed-vector versus goal-directed prediction errors with a paired Wilcoxon signed-rank test.

## SUPPLEMENTARY METHODS

### Animal preparation and chronic recordings

Animals were obtained from a local provider and maintained with food and water available ad libitum. During surgery, anesthesia was maintained by administering 15% of the initial ketamine/xylazine dose every 90 min through an intramuscular cannula. Topical 4% lidocaine was applied to wounds and pressure points. ECG was continuously monitored, and body temperature was maintained at 38–42 °C with a thermoregulated blanket (FHC).

A 3 × 3 mm craniotomy was made over the telencephalic region above the GLv. Implanted electrodes were fixed with dental acrylic and the cranial window was closed with bone wax. For awake head-fixed experiments, skull screws were implanted to permit head fixation without pressure on the head. After surgery, enrofloxacin (5 mg/kg, i.m.) and ketoprofen (3 mg/kg, i.m.) were administered daily for 5 d. Chronic recording sessions began 2 d after surgery and lasted no more than 2 h, twice daily, for 1 wk.

For optic tectum ablation, a temporal craniotomy was performed and tectal tissue was removed with a sterile bent-tip glass Pasteur pipette coupled to a vacuum trap and vacuum pump (BOECO Germany, R-300). The resulting cavity was filled with absorbable hemostat and sealed with bone wax.

### Electrical stimulation and VEP recording

Monopolar electrodes consisted of epoxy-coated tungsten rods (125 μm diameter, 20 μm exposed tip; FHC Inc.) with an impedance of approximately 1 MΩ at 1 kHz. Constant-current, electrically isolated pulse trains were delivered with an STG4002-1.6mA stimulator (Multi Channel Systems). Stimulation parameters were selected from ranges previously used to evoke orienting head and gaze movements (du Lac and Knudsen, 1990; Masino and Knudsen, 1993) and adjusted empirically at each GLv site. Current was verified from the voltage drop across a 1 MΩ resistor in series with the stimulator return lead and monitored continuously with an oscilloscope. For field recordings, the eyelid was held open with tape and VEPs (3–200 Hz) were elicited by light flashes. Signals were recorded along the dorsoventral axis of the brain and digitized at 20 kHz with a National Instruments A/D converter.

### Current source density analysis

CSD analysis requires an anatomical organization in which the recording-depth axis can be aligned with the principal axis of current flow. The GLv is well suited to this approach because its retinotopic organization extends primarily in the rostrocaudal and mediolateral planes, while retinal afferents enter ventrally, GLv-li neurons are arranged along the dorsoventral axis, and their efferents leave dorsally (Crossland and Uchwat, 1979; Guiloff et al., 1987; Vega-Zuniga et al., 2014). CSD profiles were therefore obtained along dorsoventral penetrations through the GLv (Crossland and Uchwat, 1979; Guiloff et al., 1987; Vega-Zuniga et al., 2014). After electrophysiological identification of the optic tract, the electrode was retracted 1,050 μm and VEPs were recorded at 50 μm intervals across 21 depths. At each depth, 12 LED flashes were delivered at 0.1 Hz from 2 m, and 1 s epochs beginning 400 ms before stimulus onset were averaged. Preparation stability was verified by repeat VEP recordings, and unstable preparations were excluded. CSD maps were calculated offline in IGOR Pro using the five-point formula (Nicholson and Freeman, 1975; Mitzdorf, 1985; Letelier et al., 2000). Because GLv conductivity has not been determined, CSD was expressed in mV/mm² assuming homogeneous conductivity in the tangential dimensions. The characteristic GLv profile, consisting of a sink in GLv-ne and a source in GLv-li, was used as a physiological landmark and confirmed by histological reconstruction and lesion mapping.

### Receptive-field mapping

RFs and retinal landmarks, including the pecten and optic axis, were mapped using a vertically oriented semitranslucent plastic hemisphere (57 cm diameter) centered on the right eye in conjunction with inverse retinoscopy (Nalbach et al., 1990). Eye displacement under anesthesia did not exceed 2°. The hemisphere could be swiveled into position and served as a screen representing visual space around the right eye (Letelier et al., 2004). A 4° spot of collimated light was flashed for 50 ms at locations within and outside the RF, with 12 flashes delivered at each tested position. Stimulus locations were represented in Hess coordinates projected onto a unit sphere centered on the eye. The RF center was defined as the location producing the maximal VEP, and responses at other positions were expressed as a percentage of this maximum. Responses across the visual field were fitted with a two-dimensional Gaussian. RF size was quantified from the 80% isoline of the fit and expressed as its angular radius in degrees.

### Movement recording

Animals showing poor postoperative recovery or signs of implant displacement were excluded from stimulation sessions. To reduce state-dependent variability in stimulation-evoked movements (Masino and Knudsen, 1993; Tehovnik et al., 2003), recordings were conducted in a dimly lit, partially sound-isolated room. Sessions were paused when animals appeared agitated or highly aroused and resumed after they returned to a resting state. Head-mounted devices weighed 0.6–5 g, corresponding to approximately 3–15% of head and neck weight.

Eye displacement was measured with a noninvasive infrared video camera (Arrington Research; 320 × 240 pixels; 2 g); during head-free experiments, the camera was temporarily secured to the dental-acrylic head cap with silicone adhesive. Neck contractions were measured with a magnetic-field sensor board (ZX-MAGNETIC) carrying a Hall-effect transducer (UGN3503). A 5 mm magnet was attached to the neck muscles, and the analog sensor output was digitized at 10 kHz. For the depth profiles in Fig. 2H, responses from different transects were aligned according to the corresponding CSD profile. Within each transect, voltage peaks were normalized to the maximal response and then averaged across transects for each depth.

Head movements were measured using both a high-speed camera and an IMU. A laser pointer attached to the head was aligned parallel to the eye–beak axis and projected onto the campimeter. The laser spot was recorded with an InLine IN10002GB high-speed camera (Fastec Imaging; 500 frames/s) synchronized to stimulation. The IMU consisted of a 3D accelerometer (SparkFun ADXL335) and a 3D gyroscope (SparkFun ITG-3200), mounted on two 2 × 2 cm boards and connected to an Arduino Uno sampling at 1 kHz.

### IMU calibration and head-movement analysis

The accelerometer X axis was aligned with the direction of the beak as a proxy for gaze direction. Head trajectories were reconstructed in Python from accelerometer and gyroscope signals following Borenstein et al. (2009) and Woodman (2007). The constant bias of the rate gyroscopes was estimated from the prestimulus period while the animal was stationary and subtracted. Gravity was removed from the accelerometer signals, which were transformed into global coordinates using orientation estimated with a complementary filter that low-pass filtered accelerometer measurements and high-pass filtered gyroscope measurements.

For stimulus–response parametrization, the head was returned to the same standard initial position before each trial, corresponding to the position used for IMU calibration. Head position was represented by the point at which the accelerometer X axis projected onto a fixed campimeter of 30 cm radius centered on the initial head position (pIMU). During calibration, the laser projection onto the campimeter (pLaser) provided a position estimate without accumulated inertial error. Parameters of the IMU reconstruction were adjusted until the final pIMU and pLaser positions differed by <2° (Supplementary Fig. 1B).

Movement amplitude was calculated as the great-circle distance between the initial and final pIMU positions. Latency was the interval from stimulus onset to movement onset. Direction was defined as 0° upward, 90° contraversive, ±180° downward, and −90° ipsiversive. Movement onset and offset were identified using an angular-velocity threshold of 20°/s.

### Localization of stimulation sites

GLv was targeted using the stereotaxic atlas of Karten and Hodos (1967). Visually evoked responses were monitored during electrode penetrations with an A-M Systems Model 3600 amplifier connected to a speaker and oscilloscope. CSD profiles localized GLv layers with approximately 50 μm precision.

At the end of recordings, marking lesions were produced with 7 μA positive direct current for 10 s. Electrode tracks and stimulation sites were reconstructed histologically from these lesions. In some cases, the microelectrode was additionally marked with DiI (1,1′-dioctadecyl-3,3,3′,3′-tetramethylindocarbocyanine perchlorate; DiCarlo et al., 1996).

### Neural tracer injections

Anterograde projections from GLv (Fig. 1A–E) were labeled by iontophoretic deposition of 2.5% Phaseolus vulgaris leucoagglutinin (PHA-L; Vector Laboratories) into medial GLv using glass micropipettes with 20–30 μm tips and a constant-current source (Mpodozis et al., 1996). PM-projecting GLv neurons (Fig. 1F–H) were labeled by pressure injection (PICOSPRITZER III, Parker Instruments) of 1% cholera toxin subunit B (CTB; List Biological Laboratories) into PM with a glass micropipette. PM coordinates were anterior 1.00 mm, lateral 1.0 mm, and depth 12.5 mm according to Karten and Hodos (1967). Animals survived for 5–7 d before histological processing.

### Histology

Animals were deeply anesthetized and perfused. Brains were postfixed overnight and cryoprotected in 30% sucrose for 3–4 d. Coronal sections were cut at 60 μm for lesion verification or 30 μm for tracer histology. Tracer-containing sections were immunoreacted with goat anti-CTB (1:15,000; List Biological Laboratories) or rabbit anti-PHA-L (1:10,000; Vector Laboratories) and visualized using the avidin–biotin–peroxidase method (ABC Elite kit; Vector Laboratories). Sections were mounted on gelatinized slides, briefly osmicated or counterstained with Giemsa, dehydrated, cleared, and coverslipped with Permount. Labeled neurons and axons were examined by light and dark-field microscopy and charted with Neurolucida.

### Fixed-vector versus goal-directed model comparison

To determine whether movements from different initial head positions were better described as repetitions of the same displacement or as movements toward a common endpoint, we compared fixed-vector and goal-directed models using leave-one-trial-out cross-validation (Stone, 1974). The fixed-vector and goal-directed alternatives follow the distinction used in previous microstimulation studies of gaze control (e.g., Knight and Fuchs, 2007). At each stimulation site, one trial was excluded and the remaining trials were used to predict its endpoint. The fixed-vector model applied the average movement vector of the remaining trials to the excluded trial’s starting position, whereas the goal-directed model used the average endpoint of the remaining trials as the prediction. This was repeated for every trial. Prediction error was the angular distance between predicted and observed endpoints and was summarized as RMS error across trials; the model with the lower RMS error provided the better out-of-sample fit. Movement vectors were averaged using circular statistics, endpoints as spherical centroids, and angular errors as great-circle distances.

### Statistical analysis

Data distributions were evaluated in GraphPad Prism using the Normality and Lognormality Tests. Gaussian and lognormal distributions were assessed with the D’Agostino–Pearson omnibus test and compared by relative likelihood. Outliers were assessed using the ROUT method (Q = 0.1%). Depth effects on normalized neck responses (Fig. 2H) were tested with a linear mixed-effects model with depth as a categorical fixed effect and transect as a random intercept. All depths were included, and the depth effect was evaluated by likelihood-ratio comparison with a model lacking the depth term.

Stimulus–response curves were fitted with the exponential function

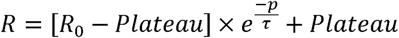

or the sigmoidal function

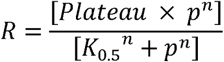

where R is the response, p is the stimulus parameter, R₀ is the response at p = 0, τ is the time constant, K₀.₅ is the value of p at half-plateau response, and n is the Hill coefficient.

Fits were weighted by 1/SD², and alternative models were compared using extra sum-of-squares F tests.

Pre- and post-ablation responses (Fig. 6) were compared using two-way ANOVA, and plateau values were compared using extra sum-of-squares F tests. RF angular radii were compared using a lognormal t test. RF–CMV associations were assessed using Pearson correlations and linear regression, with regression slopes and intercepts compared by extra sum-of-squares F tests. RMS endpoint prediction errors for the fixed-vector and goal-directed models were compared using a paired Wilcoxon signed-rank test. P < 0.05 was considered significant. Conventional analyses and curve fitting were performed in GraphPad Prism; linear mixed-effects analyses and leave-one-trial-out predictions were performed in Python.

## ACKNOWLEDGMENTS

We gratefully acknowledge Solano Henriquez and Elisa Sentis for expert technical help.

