## Supplementary material for "Retinotopic organization of head-orienting vectors in the pigeon ventral lateral geniculate nucleus"

Supplementary figure 1 (Figure 2 related)

A

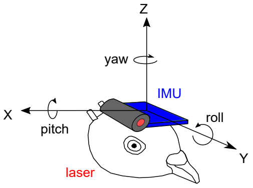


B

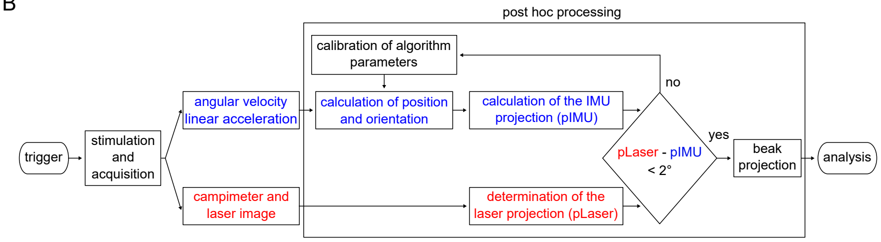

Supplementary figure 2 (Figure 6 related)

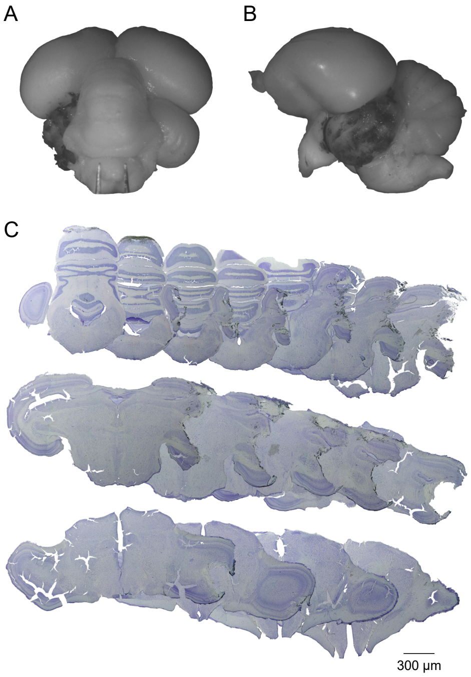

Supplementary table 1

| Parameter | Site 1a | Site 1b | Site 2 | Site 3a | Site 3b | Site 4 |
|---|---|---|---|---|---|---|
| Ipsilateral eye movement | | | | | | |
| Horizontal amplitude (°) | 5 | 2 | 4 | 3 | 3 | 6 |
| Vertical amplitude (°) | 6 | 2 | 0 | 1 | 2 | 1 |
| Threshold (µA) | 150 | 120 | 200 | 250 | 150 | 200 |
| Latency (ms) | 85 | 103 | 88 | 112 | 120 | 97 |
| Direction | nasal–ventral | nasal–dorsal | ventral | temporal–ventral | nasal–ventral | nasal–ventral |
| Contralateral eye movement | | | | | | |
| Horizontal amplitude (°) | 2 | 2 | 5 | 3 | 4 | 4 |
| Vertical amplitude (°) | 5 | 3 | 3 | 2 | 3 | 5 |
| Threshold (µA) | 200 | 120 | 150 | 200 | 180 | 200 |
| Latency (ms) | 92 | 100 | 90 | 108 | 137 | 126 |
| Direction | temporal–ventral | temporal–ventral | temporal–ventral | nasal–dorsal | nasal–ventral | temporal–dorsal |
| Head movement | | | | | | |
| Amplitude (°) | — | — | >30 | >30 | >8 | >30 |
| Threshold (µA) | — | — | 40 | 50 | 30 | 40 |
| Latency (ms) | — | — | 61 | 65 | 32 | 53 |
| Direction | — | — | contra | contra–ventral | contra–ventral | contra–ventral |
